# Site-specific RNA Functionalization via DNA-induced Structure

**DOI:** 10.1101/2020.08.06.238576

**Authors:** Lu Xiao, Maryam Habibian, Eric T. Kool

**Affiliations:** Department of Chemistry, ChEM-H Institute, and Stanford Cancer Institute, Stanford University, Stanford, CA 94305

## Abstract

Site-specific RNA functionalization is in high demand, but remains a challenge, particularly for RNAs produced by transcription rather than by total synthesis. Recent studies have described acylimidazole reagents that react in high yields at 2’-OH groups in RNAs. To date, the reactions occur stochastically at non-base-paired regions of RNA, covering much of the RNA in scattered acyl esters. Localized reactions, if possible, could prove useful in many applications, providing functional handles at specific sites and sequences of the biopolymer. Here we describe a DNA-directed strategy for *in vitro* functionalization of RNA at site-localized 2’-OH groups. The method, RNA Acylation at Induced Loops (RAIL), utilizes complementary helper DNA oligonucleotides that expose gaps or loops at selected positions while protecting the remainder in DNA-RNA duplexes. Reaction with acylimidazole reagents is then carried out, providing high yields of 2’-OH conjugation at predetermined sites. Subsequent removal of the DNA provides the RNA functionalized as desired. Experiments reveal optimal helper oligodeoxynucleotide designs and conditions for the reaction, and tests of the approach were carried out to control ribozyme activities and to label RNAs with dual-color fluorescent dyes. The RAIL approach offers a simple new strategy for site-specific labeling and controlling RNAs of any length and origin.

## Introduction

The complexity of the cellular transcriptome has raised challenges for biologists and chemists to analyze and understand the dynamics, folding and function of many individual RNA species^1^. Among the most important chemical strategies for analysis of biopolymers is the development of selective and efficient methods of chemical functionalization. Such methods enable imaging^2^, quantification^3^, structural analysis^4^, and immobilization^5^, all essential tools in biology and medicine^6^. While the development of selective chemical functionalization reactions for proteins has made strong progress over decades^7^, the development of methods for forming chemical bonds with RNAs remains at an earlier stage. To be sure, total oligonucleotide synthesis enables high levels of control over conjugate design^8^, but this technology is limited to RNAs ca. 100nt or shorter, and is inaccessible to non-specialist laboratories and commercially expensive. Among the most significant approaches developed to date for forming bonds with long or transcribed RNAs are enzymatic methods for 5’ end and 3’ end derivatization^9^, as well as enzymatic approaches applied internally in RNAs that are engineered to contain specific structural domains^6a,10^. Also investigated have been methods involving incorporation of unnatural bases via RNA polymerase^11^, or assisted either by aptamer derived from *in vitro* selection^12^ or by guide RNA-directed enzymatic modification^13^. Non-enzymatic methods are much more rare, but include 3’ end periodate oxidation followed by reaction of the resulting aldehydes^14^, which suffers issues with instability. Methods for bond formation at internal sites in transcribed RNAs remain quite limited or nonselective. DNA-templated generation of 1, N^6^-ethenoadenosine derivatives provides a way for site-specific RNA labeling^15^, but the procedure is long requiring many steps and the labeling is low-yield. Diazoketones can be used to react at random phosphates in the chain; however, at internal sites this generates RNA phosphotriester linkages that are hydrolytically unstable^16^. Photocrosslinkers such as psoralens are selective only for double-stranded regions in folded RNAs, and are used not for specific functionalization, but rather in low yields for structural mapping^17^. Alkylating agents such as dimethylsulfate that react at cytosine and adenine show only selectivity for unpaired bases over paired ones^18^, and can render the bases labile, thus losing the conjugation. As a result, this latter reaction is also used in low yields as a structure mapping strategy^19^. Finally, a number of acylating reagents that form bonds at 2’-OH groups of unpaired nucleotides have been widely used in trace yields for structural mapping^20^; however, these are typically unsuitable for high-yield functionalization due to their low solubility and very high reactivity. Importantly, no method for sequence- or site-selective reaction has been reported for any of these essentially stochastic reactions.

Recent studies have shown that increasing the solubility and lowering the reactivity of acylating agents can increase yields of 2’-OH ester formation with RNA to stoichiometric and even superstoichiometric levels^21^. A number of acylimidazole reagents have been shown to functionalize RNA at high yields, enabling efficient labeling and caging of transcribed RNAs^21d,22^. Similarly, isatoic anhydride reagents have recently been detuned and solubilized to increase functional yields for potential applications such as separation of RNA from DNA^23^. To date, however, these acylation reactions are nonselective among single-stranded regions of RNA, and so RNAs of interest contain many possible sites of reaction. This nonselective polyacylation can result, for some applications, in undesired blocking of binding, folding, and function, and places limits on control of RNA function in the development of caging strategies. Despite the increasing utility of 2’-OH acylation^24^, there remains no method for directing esterification to specific RNA sites.

Here we address this selectivity issue by developing a versatile strategy for site-localized acylation of RNAs. The method, RNA Acylation at Induced Loops (RAIL), takes advantage of the low reactivity of RNAs in double-stranded structure by use of DNA oligonucleotides to protect all but the desired reaction sites in an RNA. We show that reactive gaps or loops in the RNA can be induced by appropriately designed and inexpensive helper DNAs, resulting in high yields of acylation at predetermined sites. The sequence-selective reaction occurs with near-nucleotide resolution, and allows for subsequent RNA labeling and control. We expect that the RAIL approach will be broadly applicable to many RNA species, and should be accessible to many chemistry and biology laboratories.

## Results

### Structural design for site-specific RNA acylation

To test the possibility of site-selective RNA acylation, we constructed duplex structures containing a loop or a gap at a specific site of RNA by incorporating designed helper DNAs (Figure 1). The helper DNAs (H) were complementary to the RNA of interest (R) with one or more nucleotides missing at a predetermined position. A water-soluble nicotinyl acyl imidazole compound (NAI-N_3_), previously shown to polyacylate RNAs^21c^, was adopted as the test acylating reagent. We hypothesized that the complementary DNA could suppress RNA acylation at otherwise reactive positions by protecting the 2’-OH groups in duplex structure, which mapping studies have shown to be less reactive^4b,20a^. Conversely, designing the helper DNAs to induce loops or leave gaps would be expected to expose specific RNA 2’-OH groups to allow the acylation at designed sites. After DNA removal, the selectively functionalized RNA could be obtained and employed in further applications.

**Figure 1.**
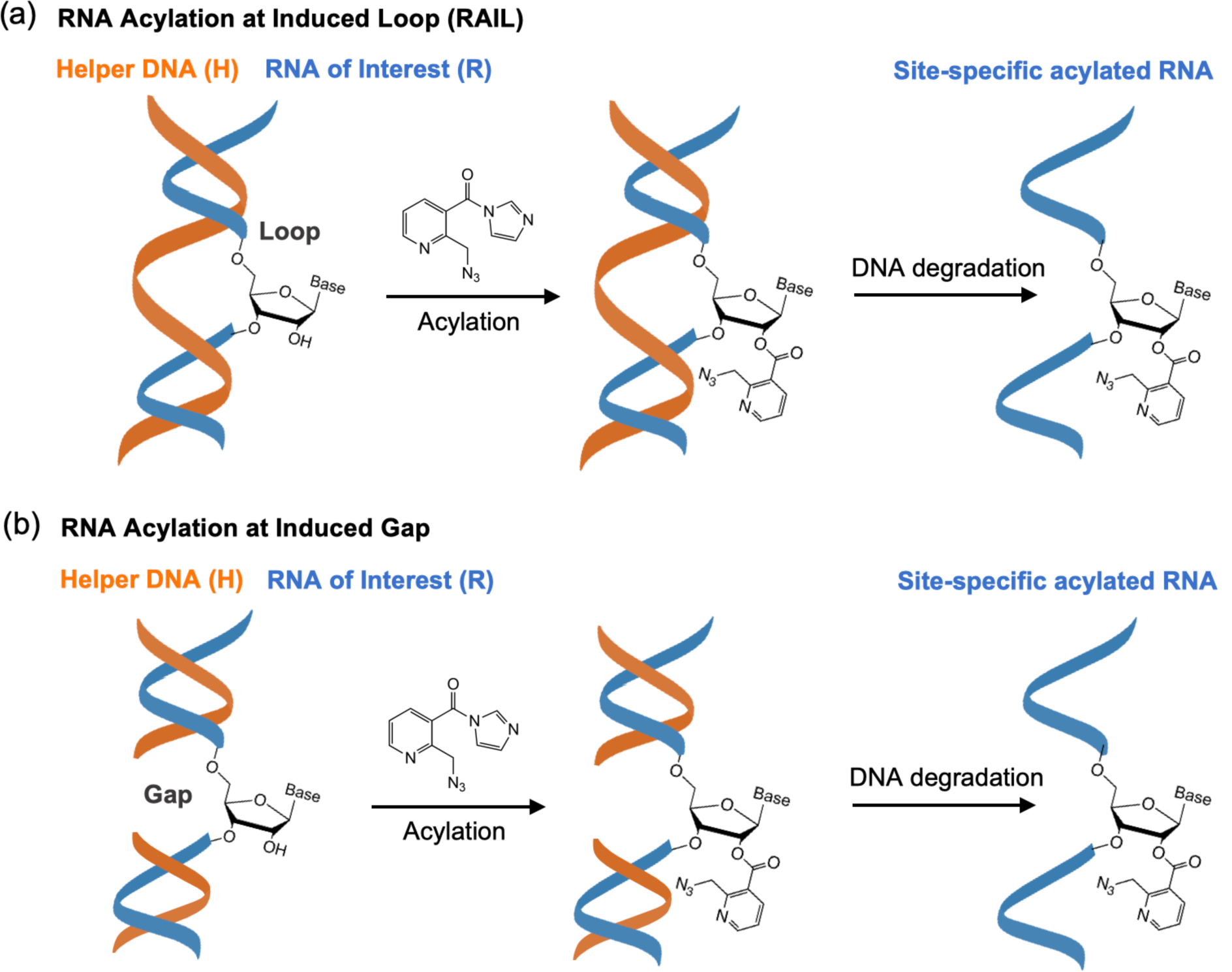
Schematic of the RAIL approach for site-specific RNA acylation. The RNA of interest (R) is incubated with one or more complementary helper DNAs (H) to protect most of the RNA, but leaving a single unpaired nucleotide in a loop or gap. The exposed 2’-OH group can then selectively react with an acylating reagent, such as the prototypical NAI-N_3_, yielding a site-specific functionalized RNA after removing the helper DNA. Note that loops and gaps larger than a single nucleotide are also possible.

### Performance of DNA-RNA duplex protection

To test our proposed strategy, we first aimed to determine the extent to which a complementary DNA can protect an RNA from 2’-OH acylation. Literature studies^4b,20^ of RNA acylation during structural mapping have established that duplex RNAs are protected relative to single-stranded sequences, but the degree of selectivity is less well characterized, particularly under high reagent concentrations necessary for stoichiometric levels of reaction. Moreover, there appears to exist no data on whether a RNA-DNA hybrid is protected similarly to purely double-stranded RNA. To test these issues, we initially studied the performance of DNA protection with a 39nt RNA model sequence derived from a biological miRNA target (Tables S1, S2). We compared the reactions of a known high-yielding (and functionalizable)^4b^ acylating agent (NAI-N_3_) with the single-stranded RNA (ssRNA) and with a pre-annealed RNA-DNA duplex by MALDI-TOF mass spectrometry to observe numbers of adducts (Figure 2a). 5 µM ssRNA or RNA-DNA duplex was reacted with 50 mM NAI-N_3_ at 37°C for 4 h in MOPS buffer, after which DNase I was added to remove the protecting DNA, and the reacted RNA was then isolated by ethanol precipitation. Importantly, we found that in comparison to ssRNA (R), which was functionalized with an average of six acyl groups (up to 25% of 2’-OH groups) under these conditions, the acylation of RNA protected by fully complementary DNA (R/H duplex) was dramatically reduced to a range of 0 to 2 groups (Figure 2b). We hypothesized that some of the observed acylation of this DNA-protected RNA might potentially occur at the ends of the RNA, since the 5’-, 2’-, and 3’-OH groups may not be efficiently protected in the blunt-ended duplex structure. To test whether the end protection could be improved, we added three-base adenine overhangs at both ends of the DNA (H_o_), expecting to stabilize the helix (R/H_o_) at the RNA ends via the stacking effects of the dangling residues^25^. Gratifyingly, this resulted in reducing acylation yields to an average of ∼0.3 acyl groups in the RNA (Figure 2b). It was not clear whether this small amount of remaining acylation occurs at the 5’ or 3’ end of the RNA, or from residual low reactivity within the duplex region. To test this, we carried out reactions with duplexes (R_p_/H_o_) in which the 2’- and 3’-OH groups at 3’ end of RNA were omitted by replacement of the terminal nucleotide with a deoxynucleoside with a 3’ phosphate group (R_p_); this resulted in almost complete loss of observed adduct peaks in the mass spectrum after reaction (Figure 2b). This establishes that the 5’-OH group is less susceptible to acylation than the hydroxyls at the 3’ end of RNA (Figure 2b, S1a). In addition, the A_3_ overhang DNA protection also worked well with a higher NAI-N_3_ concentration (Figure S1b), which can enable higher stoichiometric yields at the intended residues. Overall, the experiments revealed that a simple, unmodified overhang helper DNA can efficiently protect an RNA from highly acylating conditions, suppressing reactivity at a given 2’-OH group by at least ca. 30-fold.

**Figure 2.**
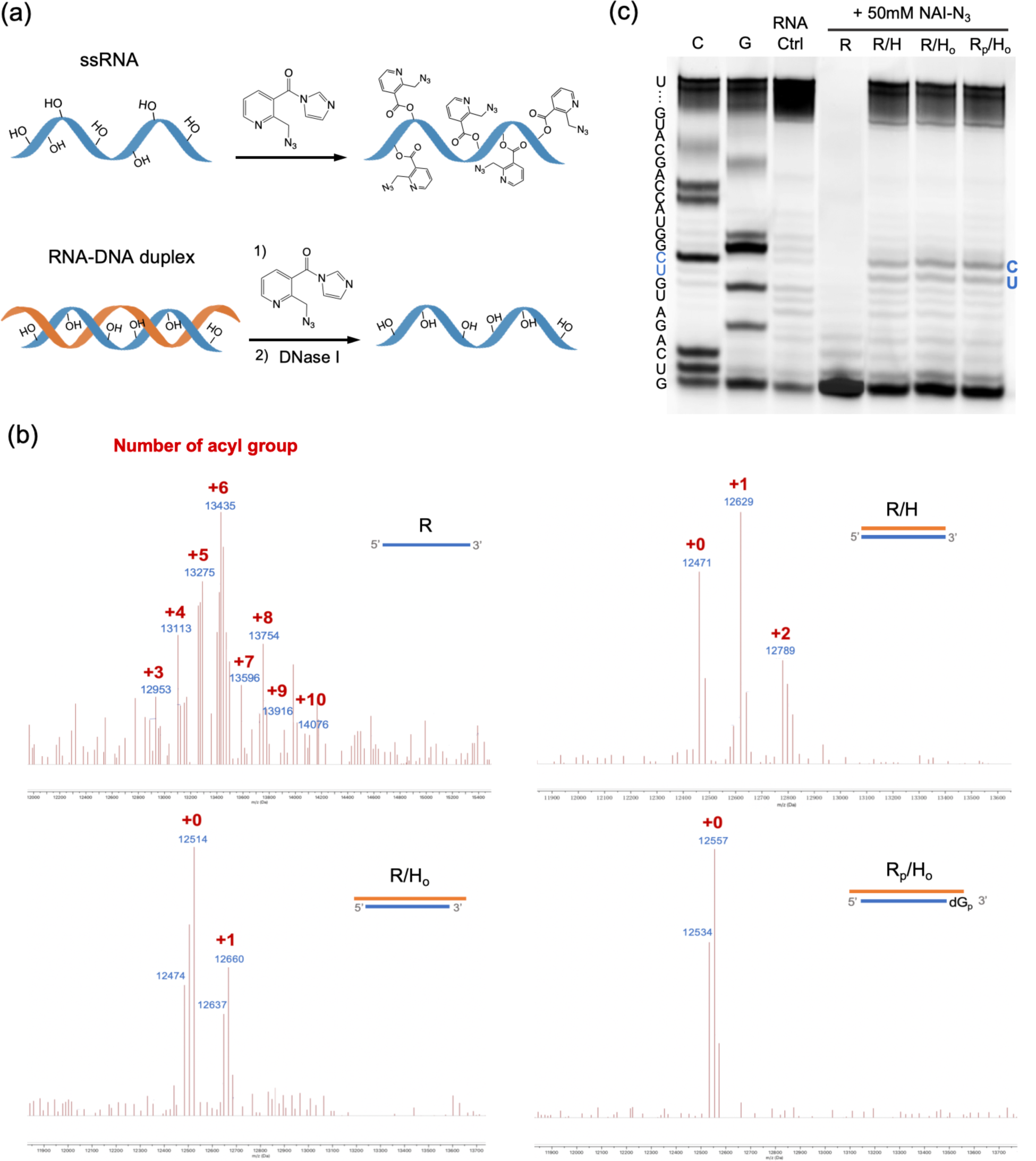
Optimizing helper DNA protection from random RNA acylation. (a) Schematic of acylation of single stranded RNA (ssRNA) and RNA-DNA duplex with a full DNA complement. (b) MALDI-TOF mass spectrum of acylated ssRNA (R), RNA protected by fully complementary DNA (R/H), RNA protected by fully complementary DNA with three deoxyadenosines overhanging at both ends (R/H_o_) and 2’-deoxy-3’-phosphate modified RNA protected by fully complementary DNA with 3a overhang (R_p_/H_o_); reacting with 50 mM NAI-N_3_ at 37°C for 4h in MOPS buffer. (c) Gel electrophoretic analysis of reverse transcriptase (RT) primer extension reveals a general lack of stops due to protected RNA, save for trace reaction at a central C, U site.

The mass spectrometry data gives useful information about yields, but not position, of acylation. To determine positions of trace acylation, we carried out studies using denaturing polyacrylamide gel electrophoresis (PAGE) after a reverse transcriptase (RT) extension of a primer hybridized to the end of the RNA. As in SHAPE structure mapping experiments^4b^, the adduct blocks RT elongation at the modified nucleotide, which results in cDNA truncation observable on the gel. The gel analysis allowed us to monitor positions of acylation at nucleotide resolution calibrated with sequencing lanes (Fig. 2c). In accordance with the mass data, most RNA-DNA duplex samples (R/H, R/H_o_, R_p_/H_o_) were successfully reverse transcribed into full-length cDNA while the unprotected and acylated ssRNA (R) was stopped at the initial stage, supporting the high performance of DNA protection in duplex (Figure 1c). The protected R/H, R/H_o_ and R_p_/H_o_ duplexes gave similarly light acylation within the body of the RNA, adding support to the prior finding that the differences in mass described above resulted primarily from small amounts of reaction at the RNA terminus. Slight bands of slightly higher trace acylation were visible at two central C, U sites compared with mock treated RNA control, which might be due to local sequence-dependent flexibility or alternative structure in the RNA-DNA duplex.

### RNA acylation at induced loops (RAIL)

Encouraged by the data showing successful DNA protection of RNA from random acylation, we next evaluated whether it is possible to restore reactivity at designed locations in the RNA by further engineering of the helper DNA. First, we tested the possibility of inducing a loop by omitting one or more nucleotides in the helper DNA, which would be expected to result in an RNA bulge loop (Figure 3a). Tests with a DNA helper designed to induce a 1nt RNA loop (R_p_/H_o_-L1) to optimize reaction conditions showed a high-yield localized RNA acylation at position “U” with 200 mM NAI-N_3_ for 4 h at 37 °C (Figure S2). To test the efficiency and site specificity of RAIL, reactions with no loop (R/H_o_ vs. R_p_/H_o_), 1nt loop (R/H_o_-L1 vs. R_p_/H_o_-L1) and 3nt loop (R/H_o_-L3 vs. R_p_/H_o_-L3) RNA-DNA duplexes were compared. We studied the prior 3’-deoxy-phosphate modified 39mer RNA (R_p_) for mass studies. The data showed a median of one adduct (range 0-2) in 1nt loop RNA (R_p_/H_o_-L1) and two adducts (range 1-3) in 3nt loop RNA (R_p_/H_o_-L3) (Figure 3b). PAGE gel analysis (Figure 3c, S3) with reverse transcriptase primer extension for the 1nt loop revealed that the darkest band appeared at G, establishing that the acylation occurs primarily at the intended bulged U residue. For the trinucleotide loop, the acylated sites were concentrated at the 3nt CUG loop sites. Interestingly, in both cases there was also a small amount of acylation at the adjacent nucleotide 5’ to the induced loop, which is explainable by added accessibility of this 2’-OH next to the bulge.

**Figure 3.**
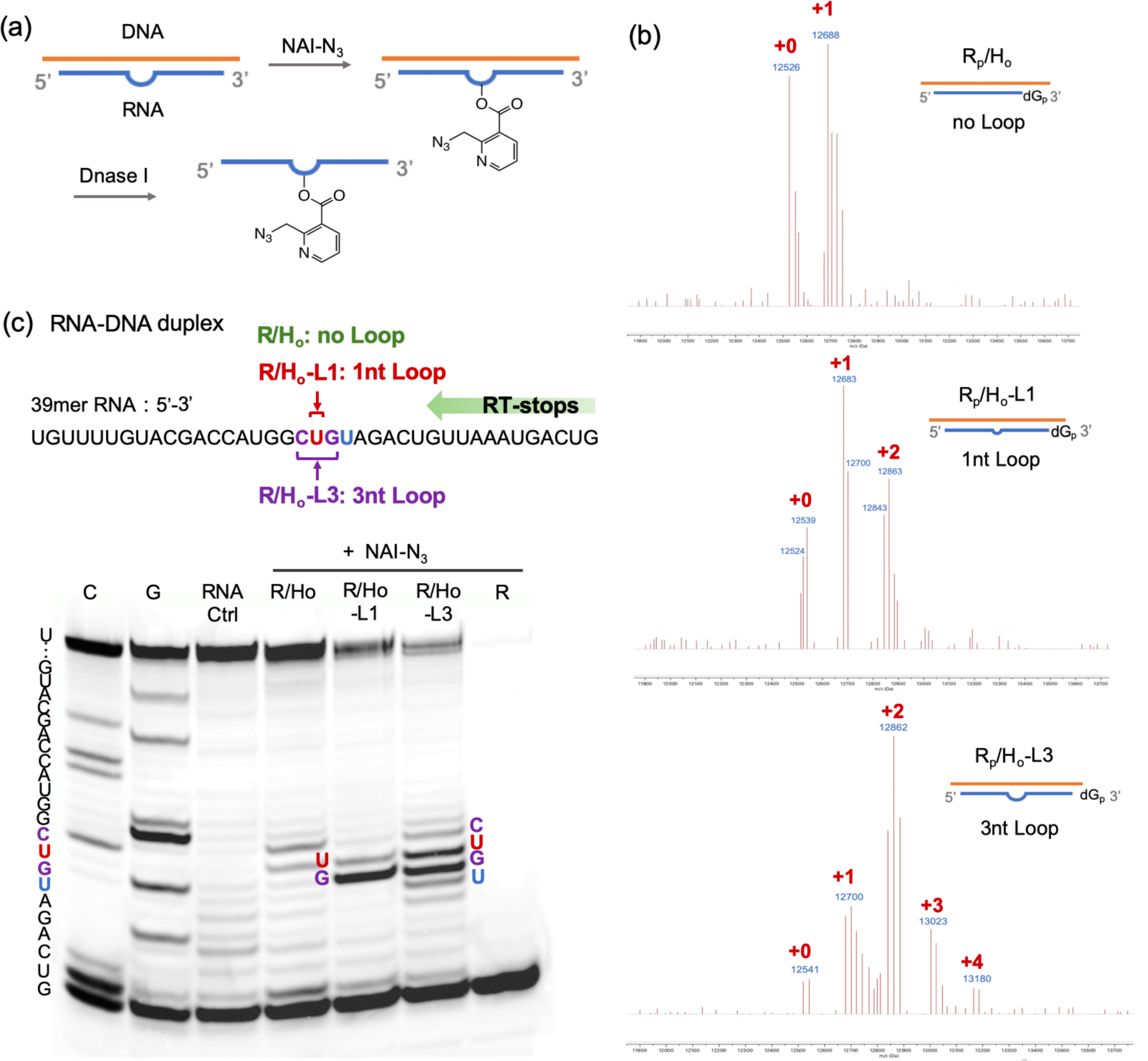
Characterization of RNA acylation at DNA-induced loops. (a) Schematic showing the process of induced loop RNA acylation. (b) MALDI-TOF mass spectra of DNA-protected RNA (R_p_/H_o_), DNA-induced 1nt loop RNA (R_p_/H_o_-L1), and DNA-induced 3nt loop RNA (R_p_/H_o_-L3), reacting with 200 mM NAI-N_3_. (c) Gel electrophoretic analysis of RT stops for the RNA samples with no loop (R/H_o_), 1nt loop (R/H_o_-L1) and 3nt loop (R/H_o_-L3) reacting with 200 mM NAI-N_3_. The sequence and sites of the bands in the gel are as shown.

### RNA acylation at an induced gap

We proceeded to study whether the localized RNA acylation could also be achieved at an induced gap position. Two strands of helper DNAs were exploited to bind adjacent to make a nicked duplex (R/H_o_-N vs. R_p_/H_o_-N), or slightly separated to render a 1nt gap at “U” (R/H_o_-G1 vs. R_p_/H_o_-G1) and 3nt gap at GCU (R/H_o_-G3 vs. R_p_/H_o_-G3) (Figure 4a). The duplexes were reacted as before with 200 mM NAI-N_3_ at 37°C in MOPS buffer. The acylation yields on RNA were found to increase from intact duplex < nick < 1nt gap < 3nt gap reactions as shown by mass spectrometry data (Figure S4). We measured the acylation positions in these cases via RT stops and again observed reaction centered at the designed gaps (Figure 4b). Notably, the nicked duplex case showed very little acylation, which supports prior observations that nicked duplexes remain largely stacked and stable^26^. Overall, we conclude that adjacently-hybridized helper DNAs are highly effective in protecting an RNA from acylation, and that a 1-nt gap left between the DNAs can result in site-selective acylation with efficiency and selectivity near that induced by a 1-nt bulge.

**Figure 4.**
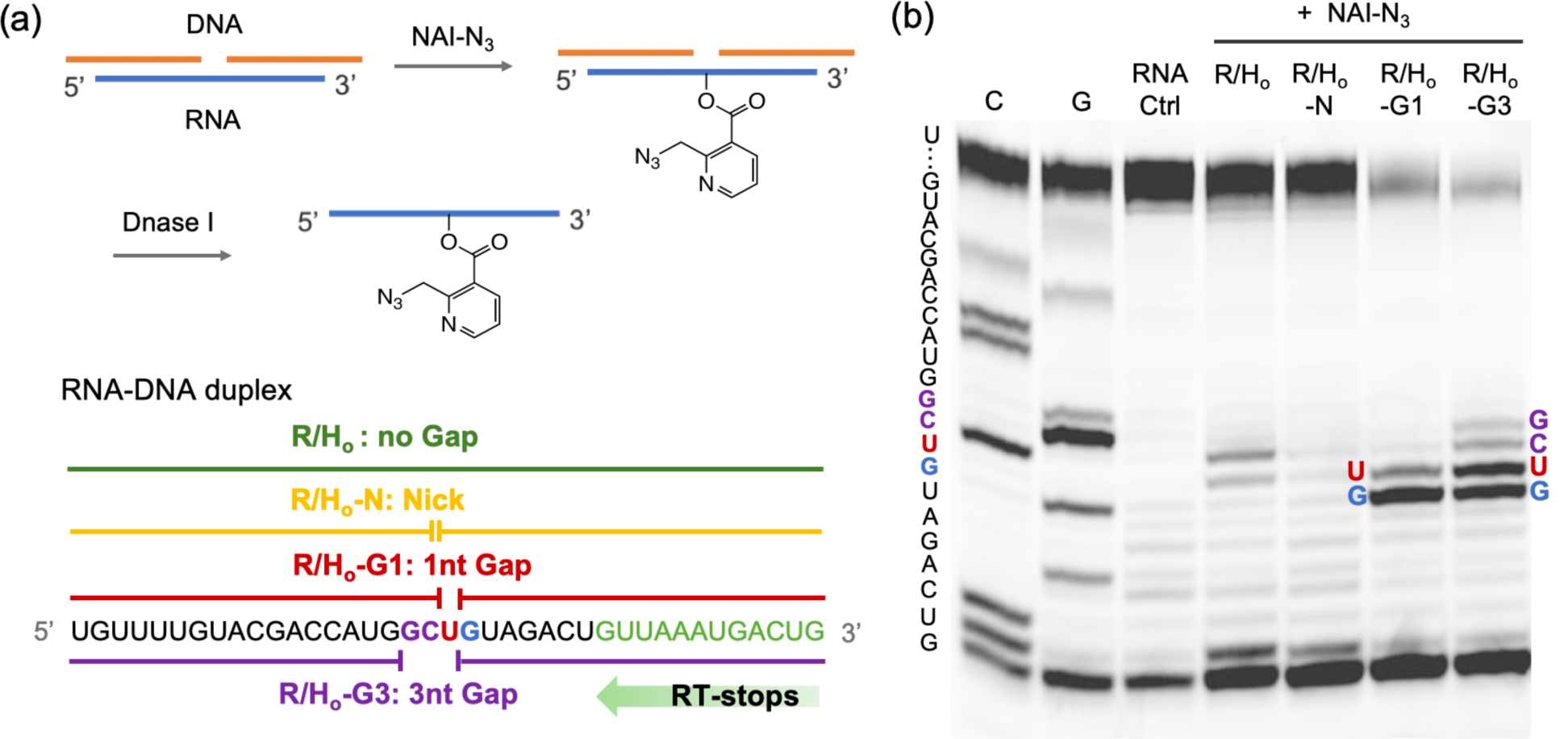
Characterization of RNA acylation at induced gaps. (a) Schematic for the steps of induced-gap RNA acylation. (b) Gel electrophoretic analysis of RT stops for RNA samples with no gap (R/H_o_), nick (R/H_o_-N), 1nt gap (R/H_o_-G1) and 3nt gap (R/H_o_-G3), reacting with 200 mM NAI-N_3_. Sequences and sites are as indicated.

### Shifting the position of acylation

After confirming that RNA acylation can be restricted to a loop or gap position, we performed additional experiments to study the generality of predetermining specific reaction positions using these methods. We therefore employed new helper DNAs to shift a putative 1nt loop or gap from U to G, C, A at progressively further locations downstream from the primer in the 39mer RNA, and measured the RT-stops after acylation, DNA removal and ethanol precipitation (Figure 5). The data show clearly that the RT stops were shifted with the moving loops or gaps, establishing programmability of the method, and establishing that loops of any of the four ribonucleotides can be acylated. Interestingly, the induced gap experiments showed poor selectivity at positions near the end, which might be expected as one helper DNA became too short to hybridize well under the reaction conditions. The induced loop cases did not show this issue, likely due to the long length of the single helper.

**Figure 5.**
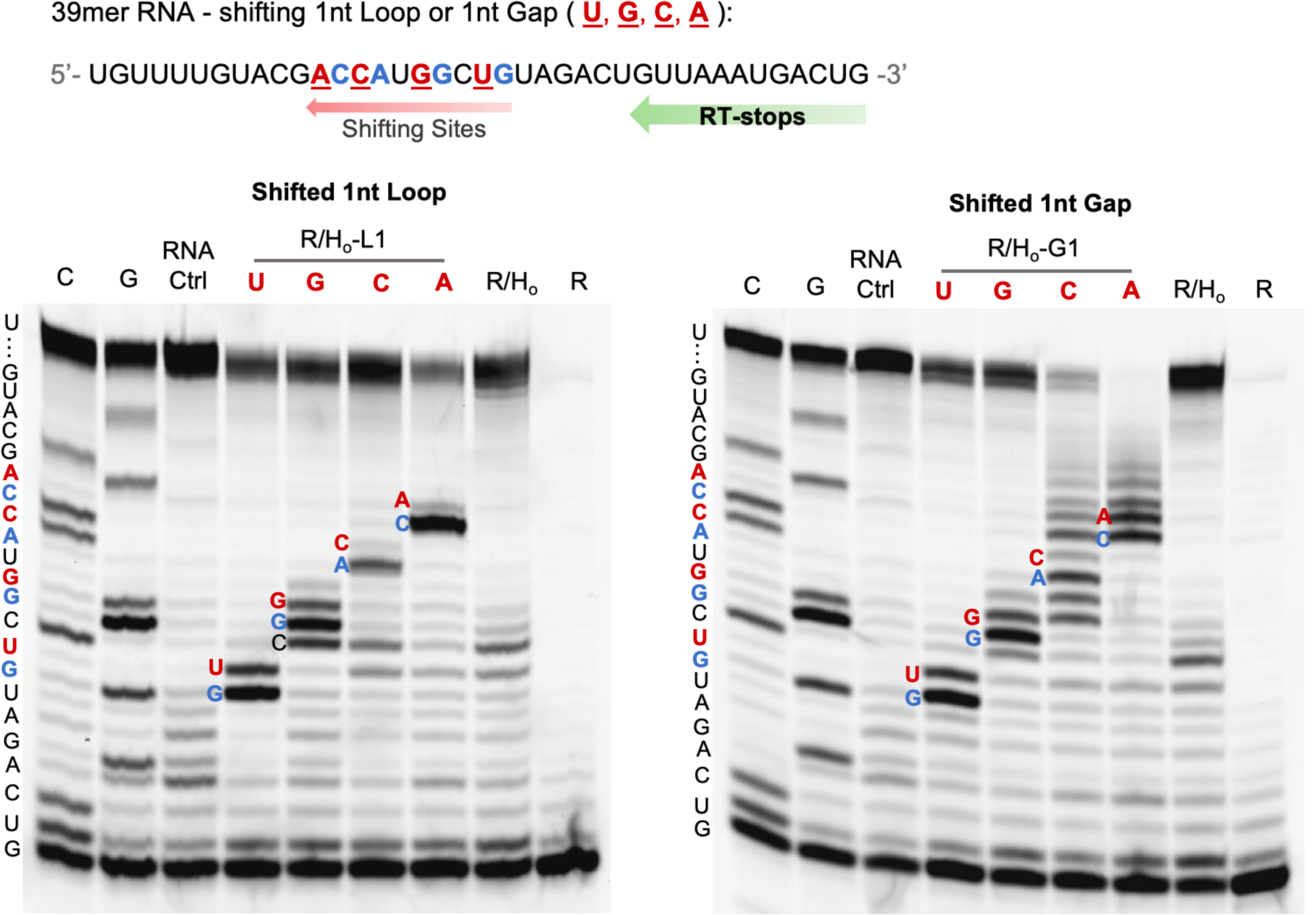
Programming local acylation sites by shifting positions of loops or gaps in a 39mer RNA target. PAGE analysis of RT-stops for the samples with 1nt loops or gaps at different positions as shown. The aligned sequences of the shifting bands in the gel are as indicated. Note that RT stops (and corresponding bands) occur at the nucleotide immediately 3’ to the reactive position.

### Site-specific control of a tandem ribozyme

Having established conditions and helper designs that yield strong protection and efficient site selectivity, we then proceeded to test the practical utility of the approach for acylation-based control of a tandem ribozyme. Although previous studies have described methods for control of a single catalytic RNA^27^, the use of a tandem ribozyme (TR) here offered the possibility to selectively and site-specifically control one RNA function relative to another within the same strand. An 81nt tandem hammerhead ribozyme with dual catalytic cores (3TR and 5TR) was produced by *in vitro* transcription and was reacted under site-selective conditions at their catalytic cores by the RAIL method under the standard conditions (Figure 6a; see SI for details). The dual catalytic activities of the TRs were measured through fluorescence enhancements by incubation with two doubly labeled reporter substrates containing a fluorophore and a quencher (3S and 5S) in 50 mM Tris pH 7.5, 10 mM MgCl_2_ (Figure 6b). The untreated TR displayed strong dual substrate cleaving activities toward 3S and 5S, giving pronounced emission at both 520 nm and 665 nm (Figure S5). The 3TR-acylated ribozyme showed a strong fluorescence at 665 nm but relatively little at 520 nm, indicating successful selective acylation at the 3TR site while conserving activity at the helper-protected site. Conversely, the 5TR acylated ribozyme using the alternative RAIL helper generated intense fluorescence at 520 nm, but weak emission at 665 nm. Analysis of the tandem ribozyme reaction rates revealed that 3TR acylation suppressed the rate of 3S cleavage by ∼5-fold and exhibited ca. 85% of the initial rate of untreated TR with the 5TR substrate; similarly, the initial cleavage rates of 5TR acylated ribozyme were decreased by 4-fold for 5S, while 3TR-ribozyme activity remained at 93% levels relative to untreated TR (Figure 6c). These results establish that RNA acylation can be locally programmed via RAIL in a multifunctional RNA, giving high yields of acylation-based suppression at the intended site, while giving little off-target acylation at the remaining sites.

**Figure 6.**
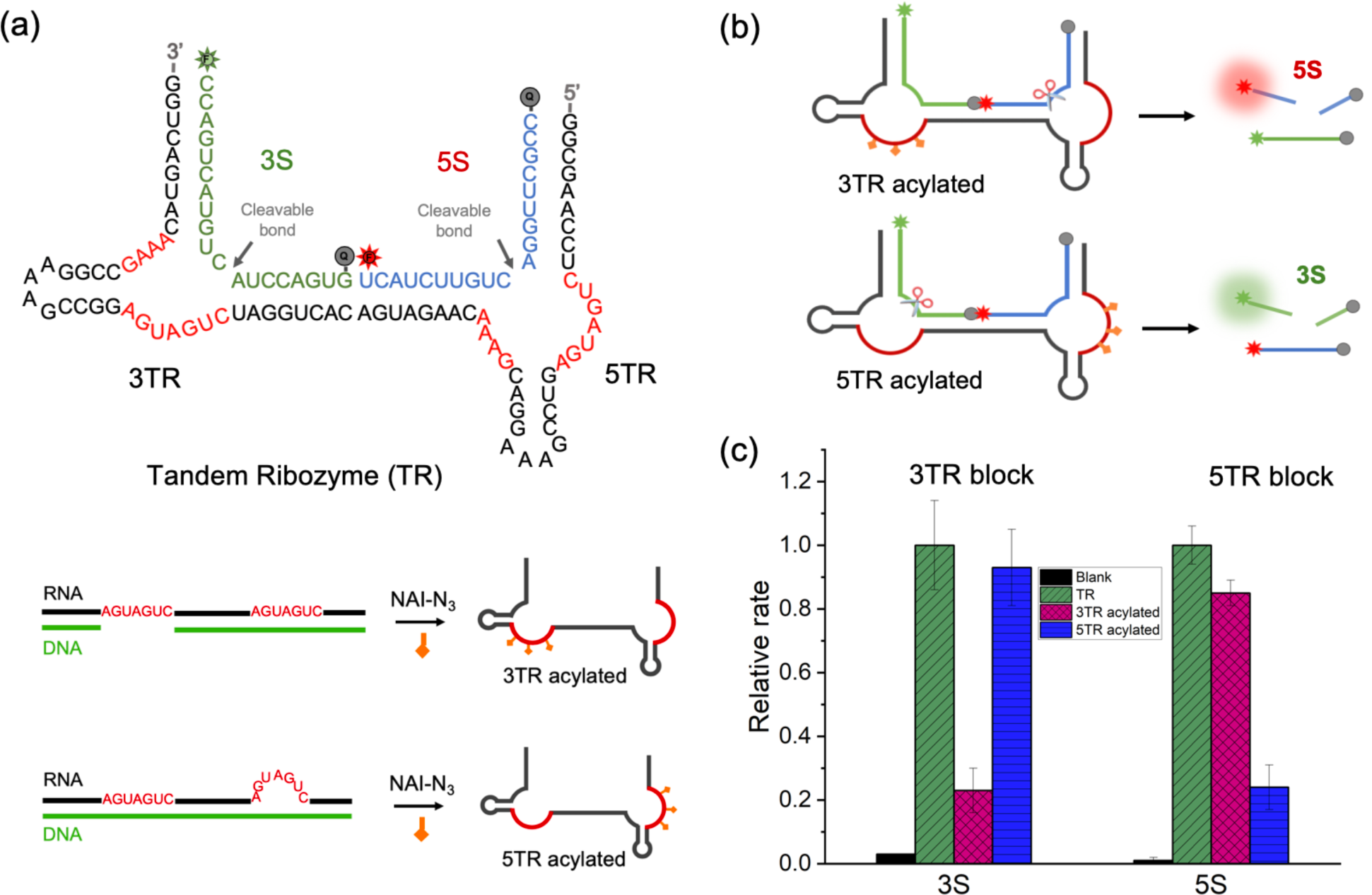
RAIL approach for the site-specific programmable control of a tandem ribozyme. (a) Sequences of a tandem ribozyme (TR) with two catalytic cores (3TR, 5TR) and two dually labeled substrates of 3S for 3TR, 5S for 5TR; Selectively acylation of TR was shown on the below via RAIL method, resulting in a 3TR acylated ribozyme and a 5TR acylated ribozyme. (b) Mechanisms of RAIL method enabled site-specific control of the tandem ribozyme. (c) Plot showing the relative initial rates of each substrate (3S and 5S) cleavage relative to that of unreacted TR (normalized to 1.0). Error bars represent the standard deviation of triplicate experiments.

### Dual labeling of small nucleolar RNA for FRET via successive RAIL labeling

The above experiments confirmed the ability of RAIL to direct site-localized suppression of one local function of a larger RNA containing multiple active sites. Next, we tested whether RAIL could be used in successive fashion on a single RNA for a dual labeling experiment. NAI-N_3_ contains both an acylimidazole group for acylation and an azide group that can react with alkynes via cycloaddition. For these experiments, we chose the 65nt small nucleolar RNA, SNORD78, which has been shown to act as an oncogene in lung cancer^28^ and is associated with hepatocellular carcinoma^29^ and prostate cancer^30^ as an enhancer for proliferation, migration and invasion. Dual labeling has been employed in synthetic DNAs and RNAs for structural studies via FRET-enabled distance measurements, but it is much more challenging to engineer a transcribed RNA to contain two labels at internal positions.

To demonstrate dual labeling of SNAORD78 RNA, we introduced fluorophore Alex488 at the 2′-OH of G14 and fluorophore TAMRA at A49 by two RAIL reactions, followed by two strain-promoted click reactions of the azide groups with DBCO modified fluorophores (Figure 7a). The labeling was carried out successively, emplacing one DNA-directed acyl group, reacting it with the first fluorescent label, then carrying out the second acylation and labeling in analogous fashion. The gel analysis confirmed the presence of both labels by the overlapping fluorescent signals of both fluorophores (Figure 7b). Labeling efficiency was sufficient to show a robust FRET signal between the dyes, and to register a drop-in energy transfer upon denaturing the structure in low-salt conditions (Figure 7c). These results suggest that the RAIL method can offer high utility in the study of RNA structure, and more generally in RNA labeling. Notably, it requires no introduction of nonnative sequences, and only knowledge of the RNA sequence is needed to carry it out.

**Figure 7.**
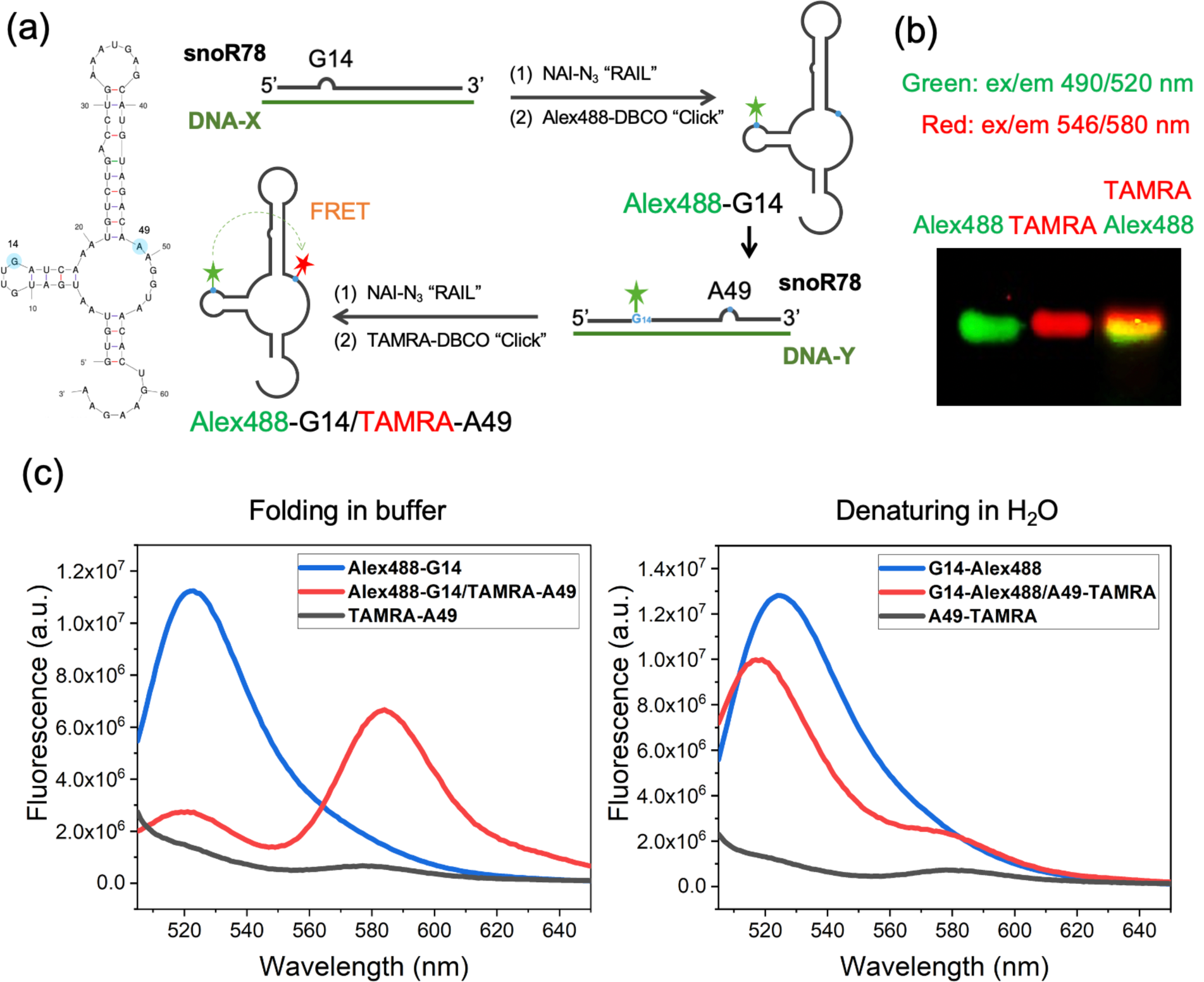
Successive RAIL approach for the dual labeling of a 65nt small nucleolar RNA (SNORD78) and observation of folding by FRET. (a) SNORD78 RNA sequence with labeling sites (G14 with Alex488; A49 with TAMRA) marked in blue. Schematic for the process of RNA dual labeling via two successive reactions. (b) Image of gel analysis of double-labeled RNA; overlay of Alex488 (green) and TAMRA (red) channels showing coincidence of the two dyes. (c) FRET signals respond to changing RNA structure in buffer versus water (fluorescence emission on donor excitation at 490 nm).

## Discussion

Our experiments introduce a novel strategy for site-directed stoichiometric acylation of RNAs produced by chemical synthesis or by transcription. The RAIL method is rapid, requiring only a few hours to yield the functionalized RNA, and its simplicity should render it accessible to chemists and biologists alike. Since acylation with NAI-N_3_ has been used both for blocking and caging^21c^ as well as click-based conjugation^4b^, the method can yield functional blocking groups, caging groups, and fluorescent labels at desired positions internally in RNA at high resolution. Our results show that it can also be performed serially on an RNA, introducing distinct labels at separate sites. The cost of the approach is low, given that DNA oligonucleotides are rapidly available at modest cost, and DNase is also widely available for removal of helper DNAs.

One limitation of the RAIL approach is the need for stable hybridization, which may restrict its use in very short RNAs (shorter than ca. 12nt) due to poor binding of helper DNAs. It is possible that helper DNAs with strongly-binding modifications may aid in addressing this issue. On the opposite end of the spectrum, it is worth considering the use of the method with DNAs longer than the available length of synthetic DNA helpers. Although synthesized DNA oligonucleotides are typically limited in length to ∼150-200nt, in principle the method could be carried out with RNAs of longer lengths by use of long enzymatically produced helper DNAs, or by tiling synthesized helper DNAs along a strand as done in our nick experiments. Future research will be needed to test this possibility. A second limitation is that placement of acyl groups at 1nt bulges or gaps by RAIL is not completely homogeneous, giving small amounts of secondary acylation (estimated at ∼20-30%; see Figure 3c) at the adjacent position. In many applications, this may not be a serious issue, in analogy to labeled antibodies where mixtures of labeled conjugates are widely useful. However, when complete RNA homogeneity is required for an experiment it is a limitation worth considering. It is possible that modification of the helper DNAs near this site may aid in suppressing this secondary acylation. Future experiments will address this issue.

## Acknowledgements

We thank the U.S. National Institutes of Health (R01GM127295 and R01GM130704) for support.

## Reference

(1) He, C., Nat. Chem. Biol. 2010, 6 (12), 863–865; Wan, Y.; Kertesz, M.; Spitale, R. C.; Segal, E.; Chang, H. Y., Nat. Rev. Genet. 2011, 12 (9), 641–655; Cech Thomas R.; Steitz, Joan A., Cell 2014, 157 (1), 77–94.

(2) Pichon, X.; Lagha, M.; Mueller, F.; Bertrand, E., Mol. Cell 2018, 71 (3), 468–480.

(3) Rabani, M.; Levin, J. Z.; Fan, L.; Adiconis, X.; Raychowdhury, R.; Garber, M.; Gnirke, A.; Nusbaum, C.; Hacohen, N.; Friedman, N.; Amit, I.; Regev, A., Nat. Biotechnol. 2011, 29 (5), 436–442.

(4) Bokinsky, G.; Rueda, D.; Misra, V. K.; Rhodes, M. M.; Gordus, A.; Babcock, H. P.; Walter, N. G.; Zhuang, X., Proc. Natl. Acad. Sci. U.S.A. 2003, 100 (16), 9302; Spitale, R. C.; Flynn, R. A.; Zhang, Q. C.; Crisalli, P.; Lee, B.; Jung, J.-W.; Kuchelmeister, H. Y.; Batista, P. J.; Torre, E. A.; Kool, E. T.; Chang, H. Y., Nature 2015, 519 (7544), 486–490; Liu, Y.; Holmstrom, E.; Yu, P.; Tan, K.; Zuo, X.; Nesbitt, D. J.; Sousa, R.; Stagno, J. R.; Wang, Y.-X., Nat. Protoc. 2018, 13 (5), 987–1005.

(5) Vioque, A.; Altman, S., Proc. Natl. Acad. Sci. U.S.A. 1986, 83 (16), 5904.

(6) Li, F.; Dong, J.; Hu, X.; Gong, W.; Li, J.; Shen, J.; Tian, H.; Wang, J., Angew. Chem. Int. Ed. 2015, 54 (15), 4597–4602; Zhang, D.; Zhou, C. Y.; Busby, K. N.; Alexander, S. C.; Devaraj, N. K., Angew. Chem. Int. Ed. 2018, 57 (11), 2822–2826.

(7) Spicer, C. D.; Davis, B. G., Nat. Commun. 2014, 5 (1), 4740; Boutureira, O.; Bernardes, G. J. L., Chem. Rev. 2015, 115 (5), 2174–2195.

(8) Allerson, C. R.; Chen, S. L.; Verdine, G. L., Journal of the American Chemical Society 1997, 119 (32), 7423–7433; Fauster, K.; Hartl, M.; Santner, T.; Aigner, M.; Kreutz, C.; Bister, K.; Ennifar, E.; Micura, R., ACS Chem. Biol. 2012, 7 (3), 581–589.

(9) England, T. E.; Uhlenbeck, O. C., Nature 1978, 275 (5680), 560–561; Paredes, E.; Das, S. R., ChemBioChem 2011, 12 (1), 125–131; Schulz, D.; Holstein, J. M.; Rentmeister, A., Angew. Chem. Int. Ed. 2013, 52 (30), 7874–7878; Samanta, B.; Horning, D. P.; Joyce, G. F., Nucleic Acids Res. 2018, 46 (17), e103–e103.

(10) McDonald, R. I.; Guilinger, J. P.; Mukherji, S.; Curtis, E. A.; Lee, W. I.; Liu, D. R., Nat. Chem. Biol. 2014, 10 (12), 1049–1054; Alexander, S. C.; Busby, K. N.; Cole, C. M.; Zhou, C. Y.; Devaraj, N. K., Journal of the American Chemical Society 2015, 137 (40), 12756–12759.

(11) Kawai, R.; Kimoto, M.; Ikeda, S.; Mitsui, T.; Endo, M.; Yokoyama, S.; Hirao, I., Journal of the American Chemical Society 2005, 127 (49), 17286–17295; Liu, Y.; Holmstrom, E.; Zhang, J.; Yu, P.; Wang, J.; Dyba, M. A.; De, C.; Ying, J.; Lockett, S.; Nesbitt, D. J.; Ferré-D’Amaré, A. R.; Sousa, R.; Stagno, J. R.; Wang, Y.-X., Nature 2015, 522 (7556), 368–372.

(12) Baum, D. A.; Silverman, S. K., Angew. Chem. Int. Ed. 2007, 46 (19), 3502–3504; Ghaem Maghami, M.; Scheitl, C. P. M.; Höbartner, C., Journal of the American Chemical Society 2019, 141 (50), 19546–19549.

(13) Tomkuvienė, M.; Clouet-d’Orval, B.; Černiauskas, I.; Weinhold, E.; Klimašauskas, S., Nucleic Acids Res. 2012, 40 (14), 6765–6773.

(14) Bellon, L.; Workman, C.; Scherrer, J.; Usman, N.; Wincott, F., Journal of the American Chemical Society 1996, 118 (15), 3771–3772; Yamamoto, J.; Ebisuda, S.; Kong, L.; Yamago, H.; Iwai, S., Chem. Lett. 2017, 46 (5), 767–770.

(15) Egloff, D.; Oleinich, I. A.; Zhao, M.; König, S. L. B.; Sigel, R. K. O.; Freisinger, E., ACS Chem. Biol. 2016, 11 (9), 2558–2567; Zhao, M.; Steffen, F. D.; Börner, R.; Schaffer, M. F.; Sigel, Roland K O.; Freisinger, E., Nucleic Acids Res. 2017, 46 (3), e13–e13.

(16) Ando, H.; Furuta, T.; Tsien, R. Y.; Okamoto, H., Nat. Genet. 2001, 28 (4), 317–325; Blidner, R. A.; Svoboda, K. R.; Hammer, R. P.; Monroe, W. T., Mol. Biosyst. 2008, 4 (5), 431–440.

(17) Lu, Z.; Zhang, Qiangfeng C.; Lee, B.; Flynn, Ryan A.; Smith, Martin A.; Robinson, James T.; Davidovich, C.; Gooding, Anne R.; Goodrich, Karen J.; Mattick, John S.; Mesirov, Jill P.; Cech, Thomas R.; Chang, Howard Y., Cell 2016, 165 (5), 1267–1279.

(18) Graveley, B. R., Mol. Cell 2016, 63 (2), 186–189.

(19) Tijerina, P.; Mohr, S.; Russell, R., Nat. Protoc. 2007, 2 (10), 2608–2623.

(20) Merino, E. J.; Wilkinson, K. A.; Coughlan, J. L.; Weeks, K. M., Journal of the American Chemical Society 2005, 127 (12), 4223–4231; Spitale, R. C.; Flynn, R. A.; Torre, E. A.; Kool, E. T.; Chang, H. Y., WIREs RNA 2014, 5 (6), 867–881.

(21) Nodin, L.; Noël, O.; Chaminade, F.; Maskri, O.; Barbier, V.; David, O.; Fossé, P.; Xie, J., Bioorg. Med. Chem. Lett. 2015, 25 (3), 566–570; Velema, W. A.; Kool, E. T., Org. Lett. 2018, 20 (20), 6587–6590; Kadina, A.; Kietrys, A. M.; Kool, E. T., Angew. Chem. Int. Ed. 2018, 57 (12), 3059–3063; Velema, W. A.; Kietrys, A. M.; Kool, E. T., Journal of the American Chemical Society 2018, 140 (10), 3491–3495; Park, H. S.; Kietrys, A. M.; Kool, E. T., Chem. Commun. 2019, 55 (35), 5135–5138; Habibian, M.; Velema, W. A.; Kietrys, A. M.; Onishi, Y.; Kool, E. T., Org. Lett. 2019, 21 (14), 5413–5416.

(22) Habibian, M.; McKinlay, C.; Blake, T. R.; Kietrys, A. M.; Waymouth, R. M.; Wender, P. A.; Kool, E. T., Chem. Sci. 2020, 11 (4), 1011–1016; Wang, S.-R.; Wu, L.-Y.; Huang, H.-Y.; Xiong, W.; Liu, J.; Wei, L.; Yin, P.; Tian, T.; Zhou, X., Nat. Commun. 2020, 11 (1), 91.

(23) Ursuegui, S.; Chivot, N.; Moutin, S.; Burr, A.; Fossey, C.; Cailly, T.; Laayoun, A.; Fabis, F.; Laurent, A., Chem. Commun. 2014, 50 (43), 5748–5751; Ursuegui, S.; Yougnia, R.; Moutin, S.; Burr, A.; Fossey, C.; Cailly, T.; Laayoun, A.; Laurent, A.; Fabis, F., Org. Biomol. Chem. 2015, 13 (12), 3625–3632.

(24) Velema, W. A.; Kool, E. T., Nat. Rev. Chem. 2020, 4 (1), 22–37.

(25) Sugimoto, N.; Kierzek, R.; Turner, D. H., Biochemistry 1987, 26 (14), 4554–4558.

(26) Protozanova, E.; Yakovchuk, P.; Frank-Kamenetskii, M. D., J. Mol. Biol. 2004, 342 (3), 775–785.

(27) Chaulk, S. G.; MacMillan, A. M., Nucleic Acids Res. 1998, 26 (13), 3173–3178; Wang, D. Y.; Lai, B. H. Y.; Sen, D., J. Mol. Biol. 2002, 318 (1), 33–43; Young, D. D.; Deiters, A., Bioorg. Med. Chem. Lett. 2006, 16 (10), 2658–2661; Savinov, A.; Block, S. M., Proc. Natl. Acad. Sci. U.S.A. 2018, 115 (47), 11976.

(28) Zheng, D.; Zhang, J.; Ni, J.; Luo, J.; Wang, J.; Tang, L.; Zhang, L.; Wang, L.; Xu, J.; Su, B.; Chen, G., J. Exp. Clin. Cancer Res. 2015, 34 (1), 49; Su, J.; Liao, J.; Gao, L.; Shen, J.; Guarnera, M. A.; Zhan, M.; Fang, H.; Stass, S. A.; Jiang, F., Oncotarget 2016, 7 (5), 5131–5142.

(29) Ma, P.; Wang, H.; Han, L.; Jing, W.; Zhou, X.; Liu, Z., Tumor Biol. 2016, 37 (12), 15753–15761.

(30) Martens-Uzunova, E. S.; Hoogstrate, Y.; Kalsbeek, A.; Pigmans, B.; den Berg, M. V.-v.; Dits, N.; Nielsen, S. J.; Baker, A.; Visakorpi, T.; Bangma, C.; Jenster, G., Oncotarget 2015, 6 (19).

