## Supplemental Information for "Site-specific RNA Functionalization via DNA-induced Structure"

Table of contents

Materials and instruments……………………………………………………………...…...S2

General procedures for site-specific acylation via RAIL method………………………...S2

Polyacrylamide gel electrophoretic analysis of reverse transcriptase stops………...…S3

Tandem ribozyme studies…………………………………………………………………S3

Dual labeling of small nucleolar RNA (SNORD78) via successive RAIL reactions…...S4

Table S1……………………………………………………………………………………..S5

Table S2……………………………………………………………………………………..S6

Figure S1……………………………………………………………………………………S7

Figure S2……………………………………………………………………………………S8

Figure S3…………………………………………………………………………………...S10

Figure S4…………………………………………………………………………...…S11

Figure S5…………………………………………………………………………………...S12

References…………………………………………………………………………………S12

### Materials and instruments

Reagents were purchased from Sigma-Aldrich unless specified otherwise. 2M NAI-$N_3$ solution in DMSO was prepared according to the previously published procedures[1]. Alex488-DBCO and TAMRA-DBCO were purchased from Click Chemistry Tools. All RNA and DNA sequences were purchased from IDT. The SuperScript™ II Reverse Transcriptase, RNaseOUT™ Recombinant Ribonuclease Inhibitor, DNase I and NTP Set (100 mM Solution) were purchased from Thermo Fisher Scientific. T7 Transcriptase was purchased from New England Biolabs (NEB).

Oligonucleotide concentrations were measured by NanoDrop One microvolume UV-Vis spectrophotometer. MALDI-TOF mass spectra were obtained on a Bruker MALDI Microflex LRF instrument using an AnchorChipTM target with the standard matrix containing 3-Hydroxypicolinic acid and diammonium citrate in TFA/acetonitrile/water. Mass spectra were analyzed with MestReNova software (v.11). MALDI-TOF mass spectra were corrected with internal standards when applicable. Fluorescence studies were performed on a Fluorolog 3-11 instrument (Jobin Yvon-SPEX). Fluorescence of the gel bands was recorded using a Typhoon 9500 laser scanner (GE Healthcare) at $\lambda_{ex}$ = 633 nm and $\lambda_{em}$ = 670 nm for Cy5, $\lambda_{ex}$ = 488 nm and $\lambda_{em}$ = 520 nm for Alex488, and $\lambda_{ex}$ = 546 nm and $\lambda_{em}$ = 580 nm for TAMARA. PAGE gel images were analyzed and quantified with ImageJ software. All quantitative experiments were performed in triplicate, and the results averaged.

### General procedures for site-specific acylation via RAIL method

50 pmol of RNA and 80 pmol of corresponding helper DNAs were heated in folding buffer containing 5 mM $MgCl_2$ to 95℃ for 2 min and cooled to room temperature to form an RNA-DNA duplex. To the annealed solutions were added 2 µL 1M NAI-$N_3$ stock in dry DMSO and 3.3 µL 3.3xMOPs buffer (333 mM MOPs, 20 mM $MgCl_2$, 333 mM NaCl, pH 7.5) to the final concentration of 200 mM NAI-$N_3$ and 5 µM RNA. The mixture was incubated at 37℃ for 4 h and then 1 µL DNase I stock was added to digest the helper DNAs at 37℃ for 30 min. 1 µL 100 mM EDTA was added prior to the degradation of DNase I at 75℃ for 10 min. After reaction, the RNA was precipitated by adding 90 µL of precipitation buffer (0.33M NaOAc, pH 5.2, glycogen 0.2 mg/mL) and 500 µL ice-cold ethanol and storing at -80 °C overnight. The resulting suspension was centrifuged at 14,800 rpm for 50 min and the supernatant was removed. The pellets were washed with 200 µL 70% EtOH, air dried, and resuspended in RNase-free water at the desired concentration. RNA concentration was determined with a Nanodrop One microvolume UV-VIS spectrophotometer. RNA samples were stored at -80 °C.

#### PAGE analysis of reverse transcriptase (RT) stops.

4 pmol RNA was mixed with 6 pmol RT Primer (Cy5-CAGTCATTTAAC) and 0.25 µL dNTP mix (10 mM each, Invitrogen), and incubated for 5 min at 65℃, then immediately chilled on ice for 2 min. Then 2 µL 5x First-Strand Buffer (Invitrogen), 1 µL 0.1 M DTT, 0.5 µL RNaseOUT and 0.25 µL Super Script II (200 U/µl, Invitrogen) were added to the final volume of 10 µL. The reaction was incubated with the following program: 25℃ for 10 min, 42℃ for 50 min, and 52℃ for 50 min. After the reaction, 10 µL loading dye (8 M Urea, 0.05% Orange G, 0.05% Bromophenol blue) was added and the mixture was denatured at 96℃ for 3 min, and loaded on a denaturing 20% polyacrylamide gel. Products were separated in a gel in 1x TBE (pH 8.3, Sigma Aldrich), 20 mA, ~2 h. The cDNA gel was visualized by fluorescence imaging (Typhoon, GE Healthcare).

#### Tandem ribozyme (TR) studies

##### TR transcription

TR was transcribed using T7 Transcriptase (NEB), according to the manufacturer's protocol, from the ordered dsDNA sequences(5'-3'):

TR DNA template Strand 1:

TAATACGACTCACTATAGGGCGAACCTCTGATGAGTCCGAAAGGACGAAACAAGATGA
CACTGGATCTGATGAGGCCGAAAGGCCGAAACATGACTGG

TR DNA template Strand 2:

CCAGTCATGTTTCGGCCTTTCGGCCTCATCAGATCCAGTGTCATCTTGTTTCGTCCTTTCG
GACTCATCAGAGGTTCGCCCTATAGTGAGTCGTATTA

##### Site-specific labeling of a tandem ribozyme (TR)

The selective acylation was performed with corresponding DNA helpers under the standard conditions as described above. For the 3TR-acylated ribozyme, we used two helper DNAs leaving a 7nt gap at the 3TR catalytic core. For the 5TR-acylated ribozyme, one helper DNA was employed to induce a 7nt bulge loop at 5TR catalytic core in the TR. The helper DNAs sequences are shown as follow(5'-3'):

helpers for 3TR acylation:

AAACCAGTCATGTTTCGGCCTTTCGGCC

ATCCAGTGTCATCTTGTTTCGTCCTTTCGGACTCATCAGAGGTTCGCCAAA

helpers for 5TR acylation:

AAACCAGTCATGTTTCGGCCTTTCGGCCTCATCAGATCCAGTGTCATCTTGTTTCGTCCTTT
CGGACAGGTTCGCCAAA

**Fluorescence assays of the tandem ribozyme (TR)**

TR RNA was first selectively acylated following the general procedures of RAIL method above. For the kinetics test, the solution contained 200 nM or 400 nM ribozyme (either TR, 3TR acylated, 5TR acylated or no ribozyme for blank) and 200 nM 3S or 400 nM 5S in 50 mM Tris and 10 mM $MgCl_2$ at pH 7.5. Sequences of 3S and 5S are given in Table S1. A time course of the cleavage reaction was immediately measured by the fluorometer at excitation/emission = 490/520 nm for FAM or 645/665 nm for Cy5, 45℃. The initial rate was calculated by the initial slope of the time course curve. The fluorescence spectra of the test solutions were measured at $\lambda_{ex}$=490 nm for FAM-3S-Q substrate and $\lambda_{ex}$=645 nm for Cy5-5S-Q substrate after incubating 30 min at 45℃.

### Dual labeling of small nucleolar RNA (SNORD78) via successive RAIL

**SNORD78 transcription**

SNORD78 was transcribed using T7 Transcriptase (NEB), according to the manufacturer's protocol, from the following dsDNA sequences:

SNORD78 DNA template Strand 1:

TAATACGACTCACTATAGGGTGTAATGATGTTGATCAAATGTCTGACCTGAAATGAGCATGT
AGACAAAGGTAACACTGAAGAA

SNORD78 DNA template Strand 2:

TTCTTCAGTGTTACCTTTGTCTACATGCTCATTTCAGGTCAGACATTTGATCAACATCATTAC
ACCCTATAGTGAGTCGTATTA

**SNORD78 dual labeling**

Helper DNA sequences (5'-3')

Helper DNA X (G14 bulge):

AAATTCTTCAGTGTTACCTTTGTCTACATGCTCATTTCAGGTCAGACATTTGATAACATCATT
ACACAAA

Helper DNA Y (A49 bulge):

AAATTCTTCAGTGTTACCTTGTCTACATGCTCATTTCAGGTCAGACATTTGATCAACATCATT
ACACAAA

100 pmol transcribed SNORD78 was first annealed to form a 1nt bulge at position "G14" with 160 pmol helper DNA (X) in folding buffer by heating to 95℃ for 2 min and cooled to room temperature,

treated with 200mM NAI-$N_3$ at 37℃ for 4h in MOPS buffer (100 mM MOPS, 6 mM $MgCl_2$, 100 mM NaCl, pH 7.5) to result in expected acylation at 2'-OH of G14. The reaction was purified by ethanol precipitation after DNase I was added to remove the helper DNA. The acylated RNA (5μM) containing the NAI-$N_3$ azide group was reacted with 25 μM Alex488-DBCO at 37℃ for 4h in 1xPBS buffer, resulting in the first labeling of Alex488-G14. The labeled RNA was purified by 10K-amicon ultra centrifugal filter. Following the same procedure, a second bulge structure at position "A49" was induced by another 160 pmol helper DNA (Y) in folding buffer after annealing. Again, after conducting the same RAIL method above, the labeled RNA with the second acylation at A49 was treated with 25 μM of the second fluorophore, TAMRA-DBCO, under the same conditions as above, to achieve dual-labeled SNORD78 RNA after final purification. 5 pmol of each labeled RNA (Alex488-G14, TAMRA-A49 or Alex488-G14/TAMRA-A49), mixed with an equal volume of loading buffer, was loaded on a denaturing 10% polyacrylamide gel and later the gel was visualized by dual channel fluorescence imaging.

**FRET measurements of SNORD78**

500 nM labeled RNAs (Alex488-G14 or Alex488-G14/TAMRA-A49) were either folded in the 1xMOPS buffer (100 mM MOPS, 6 mM $MgCl_2$, 100 mM NaCl, pH 7.5) or denatured by dissolving in water. The fluorescence spectra of the test solutions were measured at $\lambda_{ex}$=490 nm, 25℃, measuring emission from 505 nm to 650 nm.

Table S1 List of RNAs used in this work

| Name | Sequence (left to right: 5' to 3') |
|---|---|
| 39mer RNA (R) | UGUUUUGUACGACCAUGGCUGUAGACUGUUAAAUGACUG |
| 39mer 2'-deoxy-3'-phosohate RNA ($R_p$) | UGUUUUGUACGACCAUGGCUGUAGACUGUUAAAUGACUdG$_p$ |
| 3S | /56-FAM/CCAGUCAUGUCAUCCAGUG/3IABkFQ/ |
| 5S | /5Cy5/UCAUCUUGUCAGGUUCGCC/3IABkFQ/ |

Table S2 List of DNAs used in this work

| **Name** | **Sequence (left to right: 5' to 3')** |
|---|---|
| Complementary DNA for 39mer RNA/ 39mer 2'-deoxy-3'-phosohate modified RNA | |
| Comp 0 | CAGTCATTTAACAGTCTACAGCCATGGTCGTACAAAACA |
| Comp 0-3'AAA | CAGTCATTTAACAGTCTACAGCCATGGTCGTACAAAACAAAA |
| Comp 0-3'5'AAA | AAACAGTCATTTAACAGTCTACAGCCATGGTCGTACAAAACAAAA |
| Como 1-3'AAA | GCCATGGTCGTACAAAACAAAA |
| Como 2-5'AAA | AAACAGTCATTTAACAGTCTACA |
| Como 3-5'AAA | AAACAGTCATTTAACAGTCTAC |
| Como 4-3'AAA | CATGGTCGTACAAAACAAAA |
| Comp 5-3'5'AAA | AAACAGTCATTTAACAGTCTACGCCATGGTCGTACAAAACAAAA |
| Comp 6-3'5'AAA | AAACAGTCATTTAACAGTCTACCATGGTCGTACAAAACAAAA |
| mt-G-Comp1-3'AAA | ATGGTCGTACAAAACAAAA |
| mt-G-Comp3-5'AAA | AAACAGTCATTTAACAGTCTACAGC |
| mt-C-Comp1-3'AAA | GTCGTACAAAACAAAA |
| mt-C-Como 3-5'AAA | AAACAGTCATTTAACAGTCTACAGCCAT |
| mt-A-Comp1-3'AAA | CGTACAAAACAAAA |
| mt-A-Como 3-5'AAA | AAACAGTCATTTAACAGTCTACAGCCATGG |
| mt-G-Comp5-3'5'AAA: | AAACAGTCATTTAACAGTCTACAGCATGGTCGTACAAAACAAAA |
| mt-C-Comp5-3'5'AAA: | AAACAGTCATTTAACAGTCTACAGCCATGTCGTACAAAACAAAA |
| mt-A-Comp5-3'5'AAA: | AAACAGTCATTTAACAGTCTACAGCCATGGCGTACAAAACAAAA |
| Primer for RT-stops | |
| RT-primer | /Cy5/CAGTCATTTAAC |
| Complementary DNA for Tandem Ribozyme | |
| 3TR gap -comp1-3'AAA | ATCCAGTGTCATCTTGTTTCGTCCTTTCGGACTCATCAGAGGTTCGCC AAA |
| 3TR gap -comp2-5'AAA | AAACCAGTCATGTTTCGGCCTTTCGGCC |
| 5TR loop-comp3-3'5'AAA | AAACCAGTCATGTTTCGGCCTTTCGGCCTCATCAGATCCAGTGTCATCTTGTTTCGT CCTTTCGGACAGGTTCGCCAAA |
| DNA template for transcription | |
| TR template | TAATACGACTCACTATAGGGCGAACCTCTGATGAGTCCGAAAGGACGAAACAAGAT GA CACTGGATCTGATGAGGCCGAAAGGCCGAAACATGACTGG |
| SNORD78 template | TAATACGACTCACTATAGGGTGTAATGATGTTGATCAAATGTCTGACCTGAAATGAG CATGTAGACAAAGGTAACACTGAAGAA |

### Additional Figures

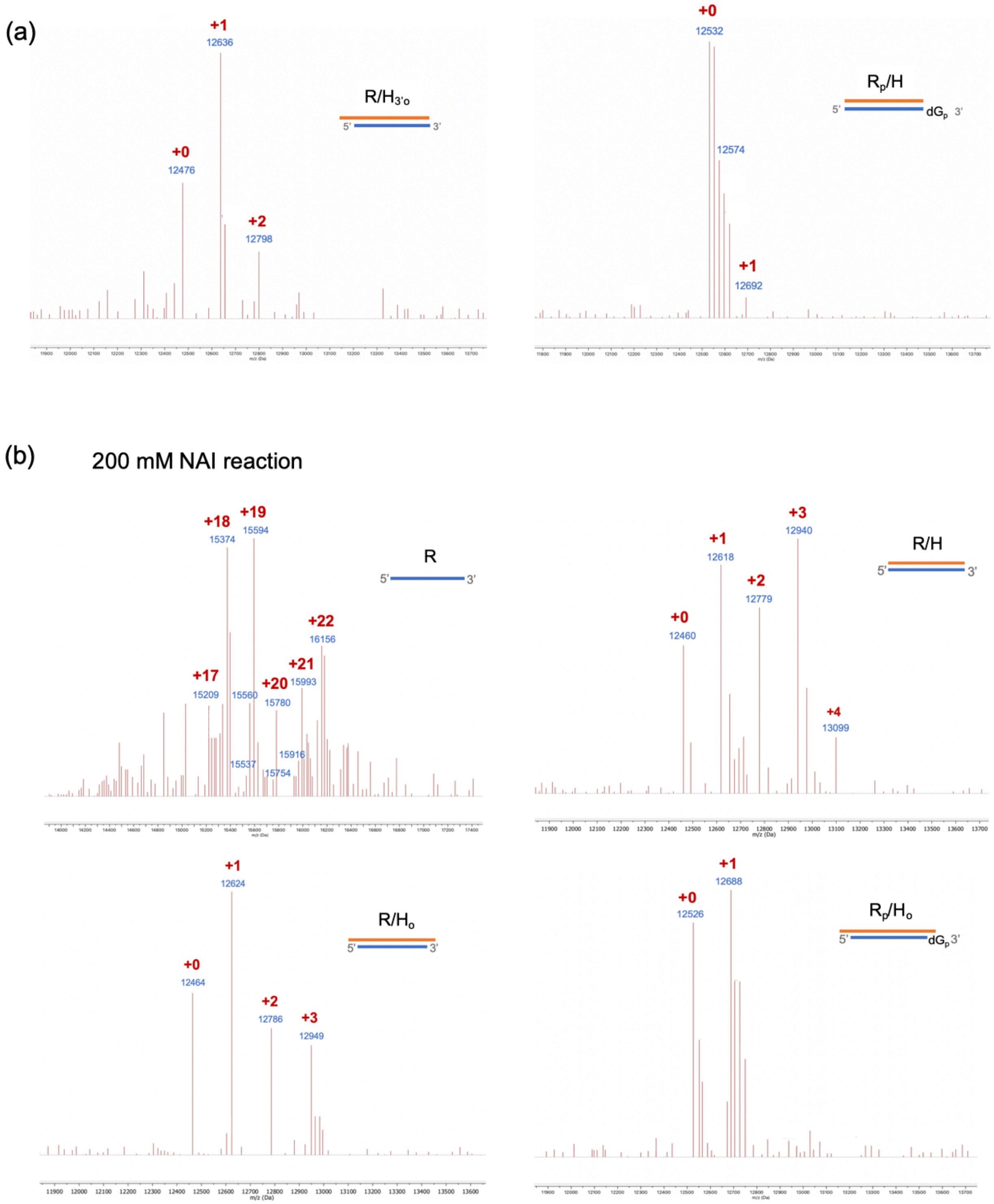


Figure S1. (a) MALDI-TOF mass spectra of RNA protected by fully complementary DNA with three deoxyadenosine overhangs at the ends ($R/H_{3'o}$) and 2'-deoxy-3'-phosphate modified RNA protected by fully complementary DNA without overhang ($R_p$/H); reacting with 50 mM NAI-$N_3$ at 37℃ for 4h in MOPS buffer. (b) MALDI-TOF mass spectrum of acylated ssRNA (R), RNA protected by fully complementary DNA (R/H), RNA protected by fully complementary DNA with three deoxyadenosines overhanging (R/$H_o$) and 3'-deoxy-phosphate modified RNA protected by fully complementary DNA with $dA_3$ overhangs ($R_p$/$H_o$), reacting with 200 mM NAI-$N_3$ at 37℃ for 4h in MOPs buffer.

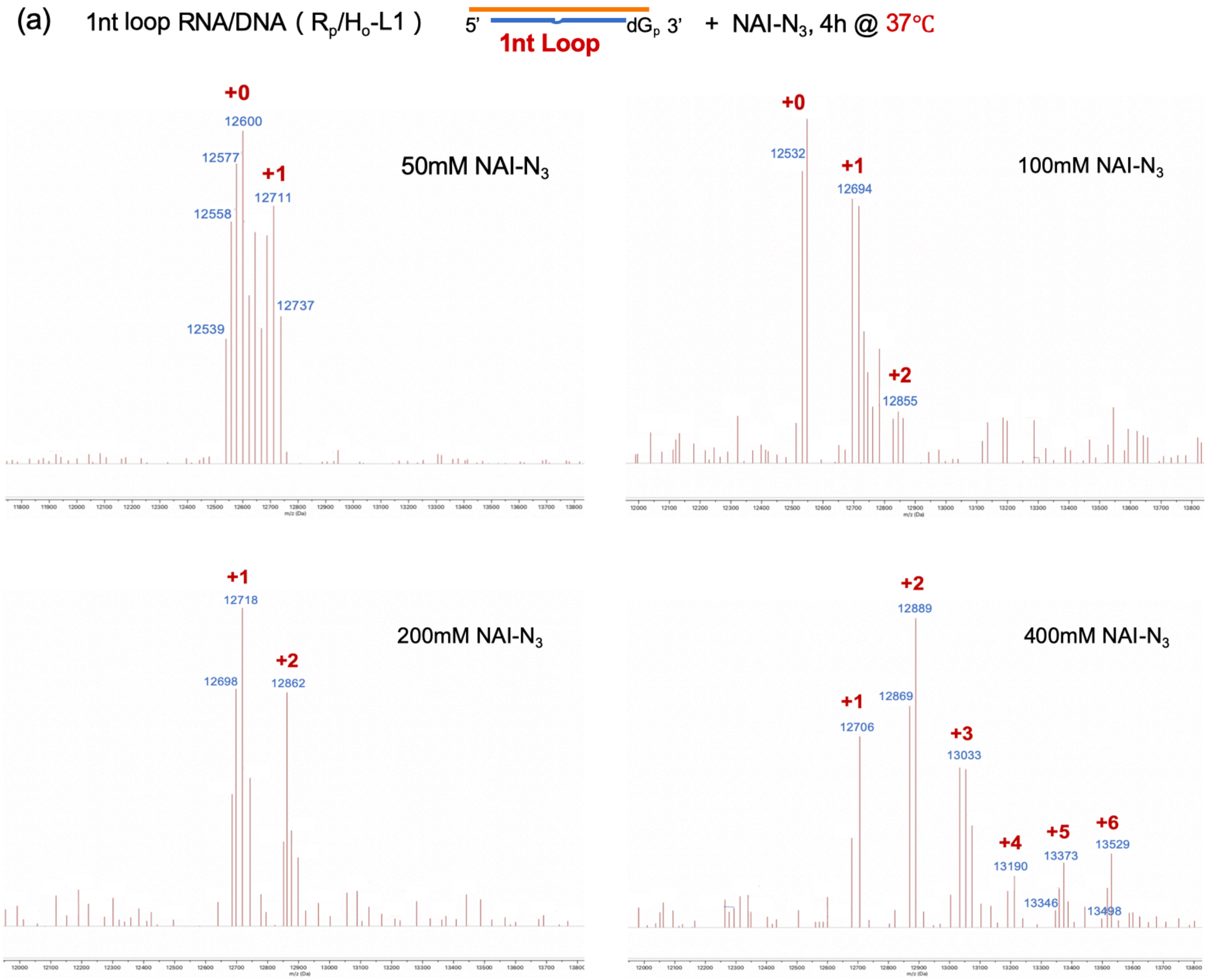

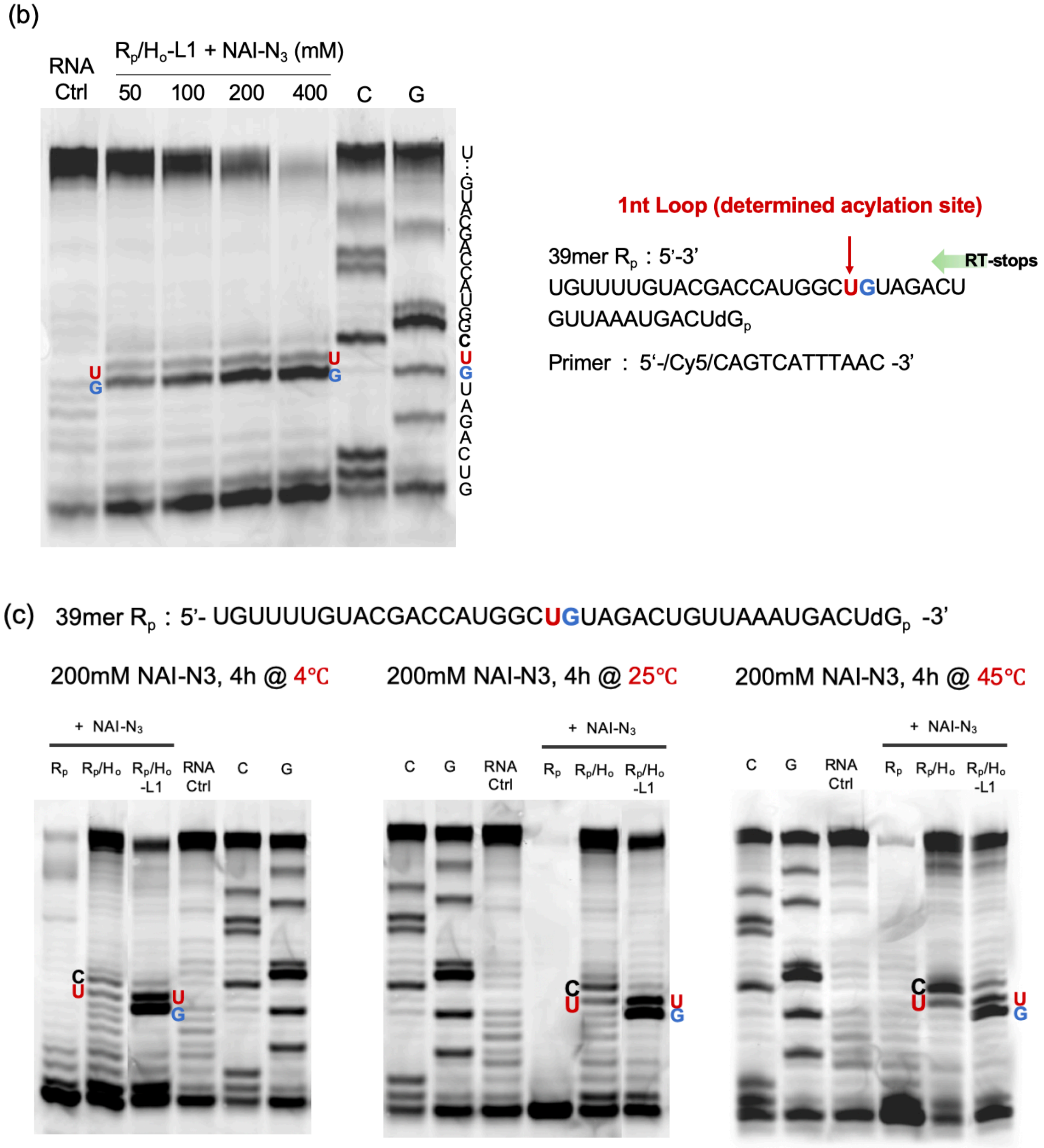


Figure S2. Optimization of RAIL conditions. (a) MALDI-TOF mass spectrum of DNA induced 1nt loop RNA samples ($R_p/H_o$-L1) reacting with different NAI-$N_3$ concentrations (50-400mM) at 37℃ for 4h in MOPs buffer. (b) PAGE analysis of the RT-stops for the $R_p/H_o$-L1 samples reacting with different NAI-$N_3$ concentrations. With increasing NAI-$N_3$ concentration, darker bands appeared at the pre-determined loop site, while full-length reverse transcribed band progressively disappears, showing an increasing yield for selective acylation. (c) PAGE analysis of the RT-stops for the

$R_p/H_o$-L1 samples reacting with 200 mM NAI-$N_3$ at different temperatures (4℃, 25℃ and 45℃) for 4h in MOPS buffer. Lower selective acylation yields (comparing with $R_p/H_o$-L1) and more random acylation (comparing with $R_p/H_o$) was observed at lower temperature, which might due to greater amounts of active residual NAI-$N_3$ remaining at the low temperature during the DNA digestion process.

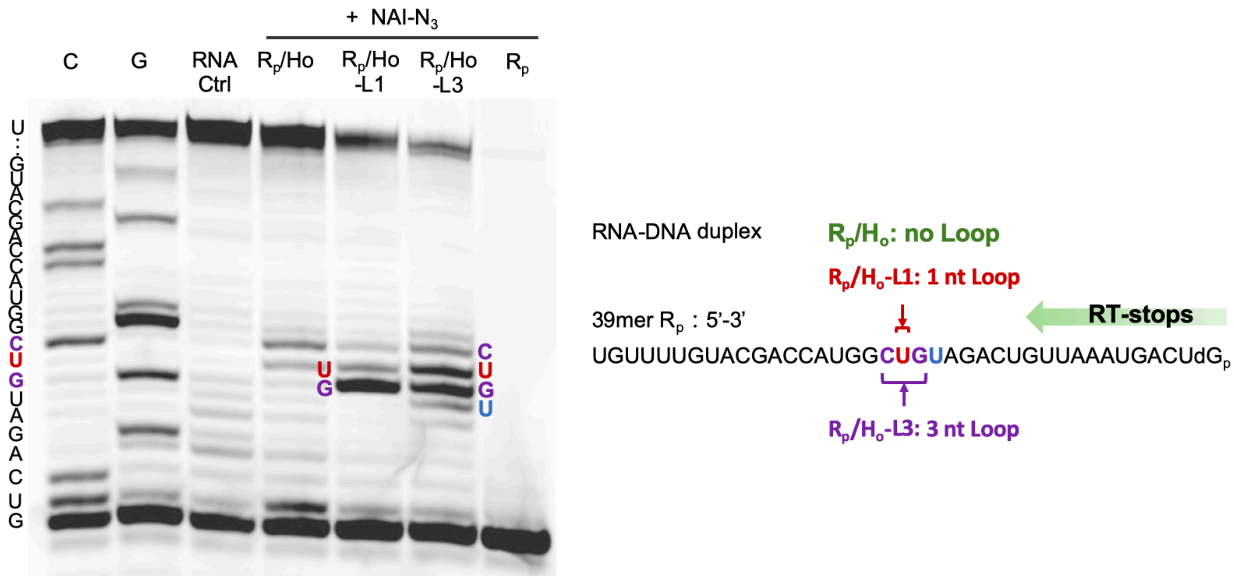


Figure S3. PAGE analysis of RT stops for the 3'-protected RNA ($R_p$) samples with no loop ($R_p/H_o$), 1nt loop ($R_p/H_o$-L1) and 3nt loop ($R_p/H_o$-L3) reacting with 200 mM NAI-$N_3$. The aligned sequences of the bands in the gel are indicated, and were identical with the natural RNA samples in figure 3c.

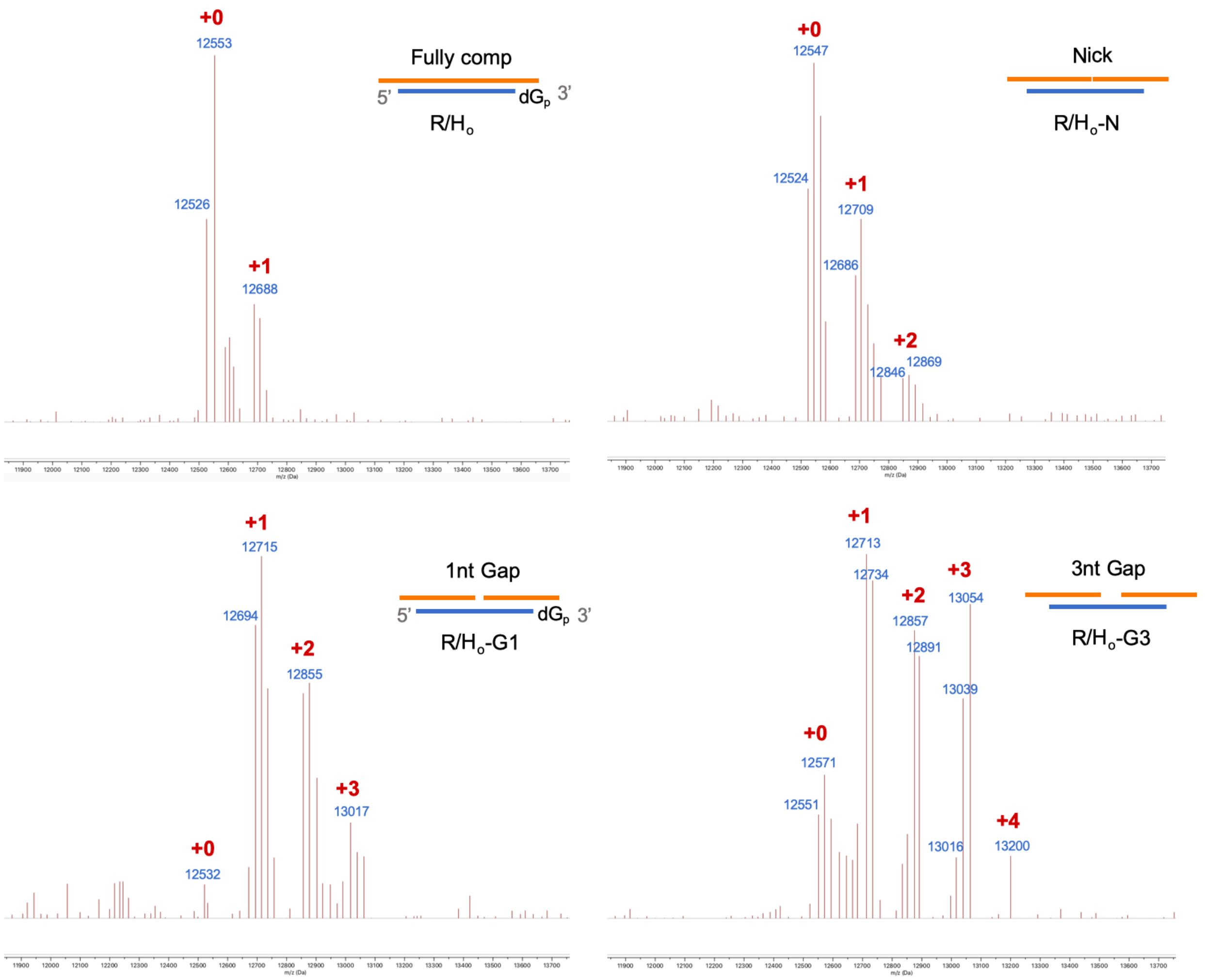


Figure S4. MALDI-TOF mass spectrum of DNA protected RNA ($R_p/H_o$), DNA protected RNA with a nick ($R_p/H_o$-N), DNA induced 1nt gap RNA ($R_p/H_o$-G1) and DNA induced 3nt gap RNA ($R_p/H_o$-G3), reacting with 200 mM NAI-$N_3$ at 37 ℃ for 4h.

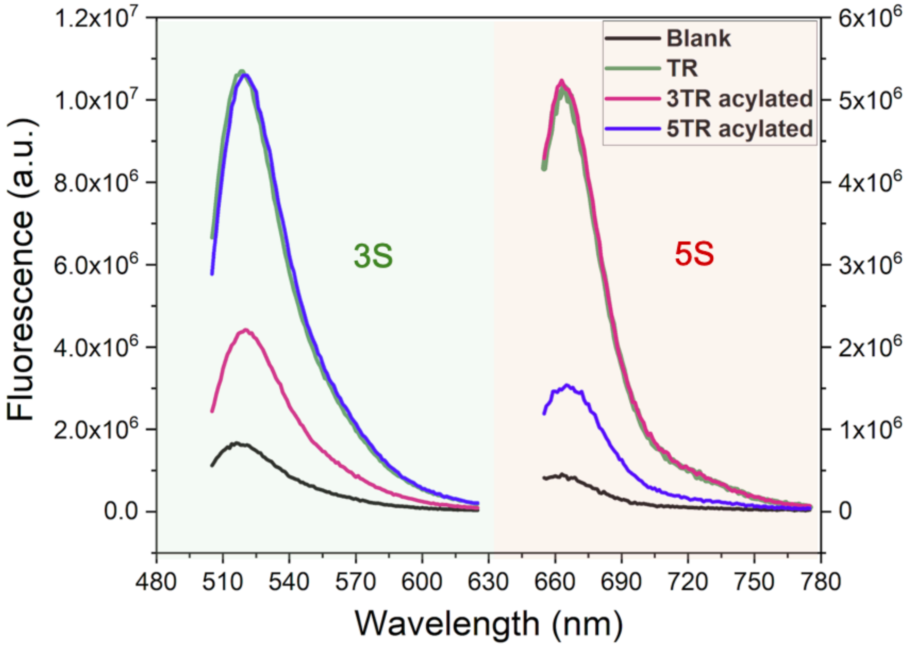


Figure S5. RAIL approach for the site-specific programmable control of a tandem ribozyme. Fluorescence emission spectra of blank (dually labeled substrate alone in buffer) (black), untreated TR (green), 3TR-acylated ribozyme (red) and 5TR-acylated ribozyme (blue) after incubation with 3S or 5S quenched fluorescent substrate RNA at 45℃ for 30 min.

References:


（1） Spitale, R. C.; Flynn, R. A.; Zhang, Q. C.; Crisalli, P.; Lee, B.; Jung, J.-W.; Kuchelmeister, H. Y.; Batista, P. J.; Torre, E. A.; Kool, E. T.; Chang, H. Y., *Nature* **2015,** *519* (7544), 486-490.
